## Supplemental Tables for "Effect of formic acid treatment on *Apis mellifera* foraging behavior as assayed using nanopore metabarcoding technologies"

**S1 Table. Times and durations of pollen collections.** Pollen was collected before Formic Acid (FA) or placebo treatment application (4/25/2022), during treatment application (4/28/2022), and at the end of treatment application (5/5/2022). Pollen was collected using pollen traps as shown in Figure 1.

| Collection 1 | Pollen Trap Down | Pollen Collected | Duration (min) |
|---|---|---|---|
| Kahlert ('K') Hives | 13:40 | 15:30 | 110 |
| HSEB ('H') Hives | 14:15 | 16:00 | 105 |
| **Collection 2** | | | |
| Kahlert ('K') Hives | 13:30 | 15:15 | 105 |
| HSEB ('H') Hives | 13:50 | 15:40 | 110 |
| **Collection 3** | | | |
| Kahlert ('K') Hives | 13:25 | 15:30 | 125 |
| HSEB ('H') Hives | 13:45 | 16:30 | 165 |

**S2 Table. Pollen DNA concentrations.** Concentrations were measured with the Nanodrop instrument (Thermo Fisher Scientific™) after extraction.

| | Hive ID | DNA Concentration |
|---|---|---|
| **Pre Treatment** | Hm1 | 256.1 |
| | Hm2 | 248.2 |
| | Hm3 | 143.6 |
| | Hw1 | 153 |
| | Ke1 | 197 |
| | Ke2 | 85.5 |
| | Ke3 | 102 |
| | Ke4 | 158.2 |
| | Km1 | 81.2 |
| | Kn1 | 98.3 |
| | Ks1 | 154.5 |
| | Ks2 | 129.5 |
| | Ks3 | 110.1 |
| **During Treatment** | Hm4 | 191.8 |
| | Ke5 | 125.5 |
| | Ke6 | 133.1 |
| | Ke7 | 91.6 |
| **Post Treatment** | Hm4 | 184.5 |
| | Hm5 | 138.4 |
| | Hm6 | 263.1 |
| | Ke8 | 111 |

| | |
|---|---|
| Ke9 | 82.6 |
| Ke10 | 73.2 |
| Km2 | 140.6 |
| Ks4 | 125.2 |
| Ks5 | 97 |
| Ks6 | 108.7 |

**S3 Table. Forward and reverse indexes.** Unique forward and reverse indexes attached to the *trnL* (UAA) intron c-d primers for the present study. The shortest hamming distance between any of the indexes is 13 base pairs and the longest is 19 base pairs.

| Forward indexes | Adenine count | Thymine count | Guanine (G) count | Cytosine (C) count | %G~C content |
|---|---|---|---|---|---|
| **agttatcgttcgaggtgttcagtg** | 4 | 9 | 8 | 3 | 45.8% |
| **atggtcggctcgatgtcatgaata** | 6 | 7 | 7 | 4 | 45.8% |
| **atgtgccgggattgaaatacgtcc** | 6 | 6 | 7 | 5 | 50% |
| **cactacctcgttcaagcttacatc** | 6 | 7 | 2 | 9 | 45.8% |
| **Reverse indexes** | | | | | |
| **ggtaaagggctatcttgtgttgcg** | 4 | 8 | 9 | 3 | 50% |
| **tcgctctgttatctaaaggccgta** | 5 | 8 | 5 | 6 | 45.8% |
| **tggtagattaactcgaacatctcc** | 7 | 7 | 4 | 6 | 41.7% |
| **tggtattgagcacctaagactgga** | 7 | 6 | 7 | 4 | 45.8% |

**S4 Table. Comparison of ONT's basecalling algorithms.** Number of reads retrieved from fast basecalling and high fidelity (SUP) basecalling for the MinION Flow Cell and Flongle runs conducted in this study. For sample information go to page 7 and 8 in the main body of the present study.

| Flow Cell | Hm1 | Hm2 | Hm3 | Hm4 | Hm5 | Hm6 | Hm7 | Kn1 | Kn2 | Ks1 | Ks2 | Ks3 | Ks4 | Ks5 | Ks6 | Ks7 | Unable to sort | Total read count |
|---|---|---|---|---|---|---|---|---|---|---|---|---|---|---|---|---|---|---|
| **Fast read count** | 306803 | 446080 | 829502 | 548950 | 298595 | 244780 | 339572 | 659166 | 524309 | 248043 | 184145 | 132440 | 220505 | 193485 | 375517 | 357576 | 755259 | 8190000 |
| **SUP read count** | 169703 | 217289 | 436745 | 291924 | 173746 | 126143 | 177041 | 340247 | 284622 | 125537 | 96003 | 62245 | 106534 | 91602 | 209989 | 171545 | 4883212 | 8317865 |

| Flongle | Hw1 | Ke1 | Ke10 | Ke2 | Ke3 | Ke4 | Ke5 | Ke6 | Ke7 | Ke8 | Ke9 | Km1 | Km2 | Unable to sort | Total read count |
|---|---|---|---|---|---|---|---|---|---|---|---|---|---|---|---|
| **Fast read count** | 3828 | 2380 | 7495 | 5684 | 5145 | 6479 | 3807 | 8450 | 7147 | 8089 | 3625 | 7981 | 7972 | 62732 | 220901 |
| **SUP read count** | 5476 | 4148 | 7619 | 6092 | 6489 | 7952 | 5316 | 8458 | 8457 | 9502 | 5014 | 9866 | 9843 | 57399 | 221101 |

**S5 Table. Demultiplexed read counts under different allowed editing distances.** Reads were demultiplexed(using minibar.py) with different maximum permitted editing distances of each index (see S3 Table), which is the individual identifying index placed at the beginning and end of each read. Read counts are from both minION flow cell and Flongle datasets, the most sorted reads occur at an editing distance of 9 ("-e 9").

| Flow Cell | 3 | 4 | 5 | 6 | 7 | 8 | 9 | 10 | 11 | 15 | 18 |
|---|---|---|---|---|---|---|---|---|---|---|---|
| **Hm1** | 106793 | 120924 | 134913 | 143395 | 150931 | 159991 | 169703 | 169750 | 175227 | 177434 | 177432 |
| **Hm2** | 129287 | 144011 | 154278 | 162804 | 173866 | 193820 | 217289 | 198132 | 198593 | 198705 | 198705 |
| **Hm3** | 291399 | 321930 | 343914 | 372226 | 395477 | 419756 | 436745 | 424781 | 424615 | 424674 | 424673 |
| **Hm4** | 172661 | 196806 | 212711 | 224877 | 243163 | 269620 | 291924 | 280743 | 280105 | 280112 | 280111 |
| **Hm5** | 105784 | 118984 | 131693 | 139197 | 148289 | 160305 | 173746 | 174380 | 176942 | 178241 | 178241 |
| **Hm6** | 72241 | 79640 | 84991 | 89339 | 96665 | 110275 | 126143 | 115470 | 115811 | 115899 | 115898 |
| **Hm7** | 108790 | 119327 | 126904 | 136534 | 147016 | 161358 | 177041 | 170000 | 169823 | 169868 | 169868 |
| **Kn1** | 202656 | 230181 | 249423 | 271581 | 293883 | 323606 | 340247 | 315990 | 316010 | 316080 | 316079 |
| **Kn2** | 153808 | 179619 | 196238 | 208484 | 232556 | 270490 | 284622 | 269749 | 268781 | 268771 | 268771 |
| **Ks1** | 72565 | 82039 | 88122 | 92730 | 100670 | 112309 | 125537 | 119345 | 118513 | 118495 | 118494 |
| **Ks2** | 54787 | 62936 | 71300 | 76079 | 81930 | 88711 | 96003 | 97006 | 99106 | 99857 | 99857 |
| **Ks3** | 29769 | 34015 | 38079 | 40700 | 45601 | 53458 | 62245 | 60634 | 61587 | 61687 | 61687 |
| **Ks4** | 60633 | 67933 | 74096 | 80536 | 87885 | 97161 | 106534 | 106361 | 108859 | 108974 | 108973 |
| **Ks5** | 50814 | 58624 | 64505 | 68355 | 74537 | 82136 | 91602 | 92514 | 91351 | 91401 | 91401 |
| **Ks6** | 127020 | 147616 | 166360 | 177585 | 189186 | 203207 | 209989 | 208872 | 212904 | 214974 | 214971 |
| **Ks7** | 96878 | 110722 | 120064 | 127317 | 137711 | 157024 | 171545 | 151233 | 151110 | 151191 | 151191 |
| **Multiple matches** | 1 | 1 | 21 | 279 | 3157 | 36338 | 353738 | 684737 | 675398 | 668895 | 669168 |

| | | | | | | | | | | | |
|---|---|---|---|---|---|---|---|---|---|---|---|
| **Unable to sort** | 6481979 | 6242557 | 6060253 | 5905847 | 5715342 | 5418300 | 4883212 | 4678168 | 4673130 | 4672607 | 4672345 |
| **Total Reads** | 8317865 | 8317865 | 8317865 | 8317865 | 8317865 | 8317865 | 8317865 | 8317865 | 8317865 | 8317865 | 8317865 |
| **FLONGLE** | **3** | **4** | **5** | **6** | **7** | **8** | **9** | **10** | | | |
| **Hw1** | 1484 | 1775 | 2003 | 2152 | 2461 | 3736 | 5476 | 9938 | | | |
| **Ke1** | 838 | 1039 | 1174 | 1264 | 1529 | 2394 | 4148 | 3412 | | | |
| **Ke10** | 2047 | 2324 | 2585 | 2796 | 3181 | 4823 | 7619 | 5224 | | | |
| **Ke2** | 1412 | 1582 | 1719 | 1805 | 2077 | 3445 | 6092 | 3573 | | | |
| **Ke3** | 1154 | 1347 | 1494 | 1644 | 2045 | 3885 | 6489 | 4307 | | | |
| **Ke4** | 1024 | 1192 | 1336 | 1541 | 2935 | 6100 | 7952 | 5164 | | | |
| **Ke5** | 1750 | 2058 | 2247 | 2357 | 2549 | 3264 | 5316 | 4952 | | | |
| **Ke6** | 2729 | 3003 | 3205 | 3358 | 3648 | 5186 | 8458 | 5572 | | | |
| **Ke7** | 2519 | 2793 | 3002 | 3245 | 3696 | 5902 | 8457 | 6590 | | | |
| **Ke8** | 1802 | 2005 | 2154 | 2371 | 4455 | 7727 | 9502 | 6948 | | | |
| **Ke9** | 1170 | 1392 | 1589 | 1695 | 1976 | 2931 | 5014 | 4294 | | | |
| **Km1** | 2052 | 2386 | 2676 | 3008 | 3651 | 6452 | 9866 | 7200 | | | |
| **Km2** | 1451 | 1687 | 1875 | 2148 | 4054 | 7641 | 9843 | 6947 | | | |
| **Multiple matches** | 1 | 1 | 4 | 13 | 142 | 4036 | 69470 | 138995 | | | |
| **Unable to sort** | 199668 | 196517 | 194038 | 191704 | 182702 | 153579 | 57399 | 7985 | | | |
| **Total Reads** | 221101 | 221101 | 221101 | 221101 | 221101 | 221101 | 221101 | 221101 | | | |

**S6 Table. DNA sequence read counts per sample before and after denoising.** Denoising consisted of removing reads shorter than 150 bp and longer than 1050 bp. Table also includes the total reads removed per sample and percentage of reads removed from the original read counts.

| Sample | Before denoising | After denoising | Reads removed | Percentage of reads removed |
|---|---|---|---|---|
| **Hw1** | 5476 | 4008 | 1468 | 26.81% |
| **Ke10** | 7619 | 5289 | 2330 | 30.58% |
| **Ke1** | 4148 | 2259 | 1889 | 45.54% |
| **Ke2** | 6092 | 3455 | 2637 | 43.29% |
| **Ke3** | 6489 | 3600 | 2889 | 44.52% |
| **Ke4** | 7952 | 4087 | 3865 | 48.60% |
| **Ke5** | 5316 | 3301 | 2015 | 37.90% |
| **Ke6** | 8458 | 5415 | 3043 | 35.98% |
| **Ke7** | 8457 | 5433 | 3024 | 35.76% |
| **Ke8** | 9502 | 5606 | 3896 | 41.00% |
| **Ke9** | 5014 | 2832 | 2182 | 43.52% |
| **Km1** | 9866 | 6139 | 3727 | 37.78% |
| **Km2** | 9843 | 5691 | 4152 | 42.18% |
| **Hm1** | 306803 | 166013 | 140790 | 45.89% |
| **Hm2** | 446080 | 213100 | 232980 | 52.23% |
| **Hm3** | 829502 | 430141 | 399361 | 48.14% |
| **Hm4** | 548950 | 282621 | 266329 | 48.52% |

| | | | | |
|---|---|---|---|---|
| **Hm5** | 298595 | 171444 | 127151 | 42.58% |
| **Hm6** | 244780 | 123358 | 121422 | 49.60% |
| **Hm7** | 339572 | 172826 | 166746 | 49.10% |
| **Kn1** | 659166 | 334069 | 325097 | 49.32% |
| **Kn2** | 524309 | 276498 | 247811 | 47.26% |
| **Ks1** | 248043 | 120380 | 127663 | 51.47% |
| **Ks2** | 184145 | 95362 | 88783 | 48.21% |
| **Ks3** | 132440 | 60706 | 71734 | 54.16% |
| **Ks4** | 220505 | 103765 | 116740 | 52.94% |
| **Ks5** | 193485 | 88610 | 104875 | 54.20% |
| **Ks6** | 375517 | 204878 | 170639 | 45.44% |
| **Ks7** | 357576 | 167046 | 190530 | 53.28% |
| **Total** | 5909468 | 3010817 | 2898651 | 49.05% |

**S7 Table. Plant genera identified from DNA sequencing (after denoising) and the corresponding family name.**

| Family | Genus |
|---|---|
| Adoxaceae | *Viburnum* |
| Amaryllidaceae | *Narcissus* |
| | *Nothoscordum* |
| Asparagaceae | *Barnardia* |
| Asteraceae | *Taraxacum* |
| | *Xanthium* |
| Berberidaceae | *Berberis* |
| | *Ehretia* |
| Fagaceae | *Fagus* |
| | *Cercis* |
| Platanaceae | *Platanus* |
| Rosaceae | *Chaenomeles* |
| | *Cotoneaster* |
| | *Cydonia* |
| | *Eriobotrya* |
| | *Malus* |

| | |
|---|---|
| | *Photinia* |
| | *Prunus* |
| | *Pyrus* |
| | *Rhaphiolepis* |
| | *Sorbus* |
| | *Stranvaesia* |
| | *Torminalis* |
| | *Vauquelinia* |
| Sapindaceae | *Acer* |
| | *Aesculus* |

**S8 Table. Pairwise PERMANOVA comparing relative read abundance data between treatments.** Significant results are highlighted in bold. Analyses were calculated using the "pairwise.perm.manova" function from the RVAideMemoire package in R and performed on Hellinger-transformed, relative read abundance data.

| | **During-Control** | **During-Formic Acid** | **Post-Control** | **Post-FormicAcid** | **Pre-Control** |
|---|---|---|---|---|---|
| **During-Formic Acid** | 0.890 | - | - | - | - |
| **Post-Control** | 0.890 | 0.243 | - | - | - |
| **Post-Formic Acid** | 0.890 | 0.890 | 0.078 | - | - |
| **Pre-Control** | 0.890 | 0.273 | 0.084 | 0.120 | - |
| **Pre-Formic Acid** | 0.408 | 0.099 | 0.042 | 0.030 | 0.264 |
| P-value adjustment method: holm | 999 permutations | | | | |

**S9 Table. Pairwise PERMANOVA comparing relative read abundance data between hive locations.** Significant results are highlighted in bold. Analyses were calculated using the "pairwise.perm.manova" function from the RVAideMemoire package in R and performed on Hellinger-transformed, relative read abundance data. For sample information see page 7 in the main body of the present study.

| | **Hm** | **Hw** | **Ke** | **Km** | **Kn** |
|---|---|---|---|---|---|
| **Hw** | 1 | - | - | - | - |
| **Ke** | 0.015 | 1 | - | - | - |
| **Km** | 1 | 1 | 1 | - | - |
| **Kn** | 0.33 | 1 | 0.657 | 1 | - |
| **Ks** | 0.052 | 1 | 0.015 | 0.33 | 0.252 |
| P-value adjustment method: holm | 999 permutations | | | | |

**S10 Table. Shannon Diversity Index summary statistics.** All relevant statistics calculated from DNA reads of plant genera collected by honeybees under different treatment conditions and locations.

| Treatment | n | mean value | median value | sd value | min value | max value | IQR value |
|---|---|---|---|---|---|---|---|
| **Pre-Control** | 5 | 1.77281218 | 1.770549503 | 0.02480089107 | 1.734333393 | 1.797386705 | 0.02190925803 |
| **Pre-Formic Acid** | 8 | 1.761852706 | 1.759693028 | 0.04497754552 | 1.685500595 | 1.829252644 | 0.05077544156 |
| **During-Control** | 2 | 1.70460138 | 1.70460138 | 0.006621382391 | 1.699919355 | 1.709283404 | 0.00468202439 |
| **During-Formic Acid** | 4 | 1.615196014 | 1.562807317 | 0.1222971021 | 1.537568329 | 1.797601095 | 0.07466669956 |
| **Post-Control** | 4 | 1.884239262 | 1.874698514 | 0.06060441889 | 1.82299984 | 1.964560177 | 0.06211617079 |
| **Post-Formic Acid** | 6 | 1.806436071 | 1.794020121 | 0.0735537505 | 1.70480044 | 1.896656522 | 0.09615493478 |
| **Location** | | | | | | | |
| **Hm** | 7 | 1.816592309 | 1.797386705 | 0.09322908504 | 1.699919355 | 1.964560177 | 0.1222570666 |
| **Hw** | 1 | 1.804681624 | 1.804681624 | NA | 1.804681624 | 1.804681624 | 0 |
| **Ke** | 10 | 1.698129969 | 1.735883603 | 0.1047503015 | 1.537568329 | 1.829252644 | 0.1607663162 |
| **Km** | 2 | 1.79647043 | 1.79647043 | 0.03751825166 | 1.76994102 | 1.82299984 | 0.02652941017 |
| **Kn** | 2 | 1.750566841 | 1.750566841 | 0.05838359649 | 1.709283404 | 1.791850278 | 0.04128343699 |
| **Ks** | 7 | 1.80117726 | 1.797601095 | 0.07305409899 | 1.685500595 | 1.896656522 | 0.07463839319 |

**S11 Table. Richness summary statistics.** All relevant statistics calculated from DNA reads of plant genera collected by honeybees under different treatment conditions and locations.

| TREATMENT | n | mean value | median value | sd value | min value | max value | IQR value |
|---|---|---|---|---|---|---|---|
| **During-Control** | 2 | 38 | 38 | 4.242640687 | 35 | 41 | 3 |
| **During-Formic Acid** | 4 | 34.25 | 33.5 | 1.892969449 | 33 | 37 | 1.75 |
| **Post-Control** | 4 | 43.25 | 43 | 3.774917218 | 39 | 48 | 3.75 |
| **Post-Formic Acid** | 6 | 37.33333333 | 36.5 | 1.751190072 | 36 | 40 | 2.5 |
| **Pre-Control** | 5 | 36.8 | 36 | 1.095445115 | 36 | 38 | 2 |
| **Pre-Formic Acid** | 8 | 37.75 | 36 | 4.832922807 | 33 | 47 | 4.5 |
| **LOCATION** | | | | | | | |
| **Hm** | 7 | 40.28571429 | 39 | 4.423960733 | 36 | 48 | 5.5 |
| **Hw** | 1 | 47 | 47 | NA | 47 | 47 | 0 |
| **Ke** | 10 | 36.2 | 35 | 3.489667288 | 33 | 43 | 5.25 |
| **Km** | 2 | 40 | 40 | 2.828427125 | 38 | 42 | 2 |
| **Kn** | 2 | 35.5 | 35.5 | 0.7071067812 | 35 | 36 | 0.5 |
| **Ks** | 7 | 36.28571429 | 36 | 0.9511897312 | 35 | 38 | 0.5 |
